# Visualizing active viral infection reveals diverse cell fates in synchronized algal bloom demise

**DOI:** 10.1101/2020.06.28.176719

**Authors:** Flora Vincent, Uri Sheyn, Ziv Porat, Assaf Vardi

## Abstract

Marine viruses are considered as major evolutionary and biogeochemical drivers of microbial life, through metabolic reprogramming of their host and cell lysis that modulates nutrient cycling^1^, primary production and carbon export in the oceans^2^. Despite the fact that viruses are the most abundant biological entities in the marine environment, we still lack mechanistic and quantitative approaches to assess their impact on the marine food webs. Here, we provide the first quantification of active viral infection, during bloom succession of the cosmopolitan coccolithophore *Emiliania huxleyi*, by subcellular visualization of both virus and host transcripts on a single cell resolution across thousands of cells. Using this novel method, that we coined Virocell-FISH, we revealed that distinct transcriptional states co-exist during the infection dynamics, and that viral infection reached only a quarter of the *E. huxleyi* population although the bloom demised in a synchronized manner. Through a detailed laboratory time-course infection of *E. huxleyi* by its lytic large virus EhV, we quantitatively show that metabolically active infected cells chronically release viral particles, and that viral-induced lysis is not systematically accompanied by virion increase, thus challenging major assumptions regarding the life cycle of giant lytic viruses. Using Virocell-FISH, we could further assess in a new resolution, the level of viral infection in cell aggregates, a key ecosystem process that can facilitate carbon export to the deep ocean^3^. We project that our approach can be applied to diverse marine microbial systems, opening a mechanistic dimension to the study of host-pathogen interactions in the ocean.

**One Sentence Summary:** Quantifying active viral infection in algal blooms

## Main Text

Microbes play a crucial role in shaping the marine ecosystem and its biogeochemical cycles^4^. In particular, abundant photosynthetic micro-eukaryotes and cyanobacteria (phytoplankton) contribute to half of the total primary production on Earth and form the basis of the marine food web ^5–7^. Phytoplankton blooms can undergo synchronized demise following infection by viruses that can reach 10^8^ viruses per L of seawater^3,8^ and are therefore hypothesized to control the fate of phytoplankton blooms and the subsequent massive release of organic matter that fuels the microbial life via the viral shunt^9^. One of the main challenges in aquatic virology is to quantify how viruses, through control of the host metabolism and abundance, remodel nutrient fluxes within major biogeochemical cycles and impact the microbial community^10,11^. This challenge meets the critical need to quantify active viral infection within host cells (virocell), and to assess the magnitude of viral-induced mortality within a complex microbial consortium in the marine environment.

Approaches such as bulk RNA-seq^12,13^, ddPCR^14^, polony^15^, viral-BONCAT^16^, phageFISH^17^ and single cell genomics^18,19^ have considerably expanded the toolbox of viral ecology and provided insights into viral diversity and viral-encoded auxiliary metabolic genes that remodel the host metabolism^20^. However, they do not report active viral infection of individual cells thereafter named “virocells”, that can be defined as “the living form of the virus in its intra- cellular form”^21^. Recent advances in single cell transcriptomics have greatly improved our understanding of host-virus interactions by uncovering high heterogeneity in the host metabolic states and viral program dynamics within individual virocells^22–24^. However, these methods lack the ability to visualize infection on the subcellular level that can provide fresh insights into the complex life cycle of viruses in their host cells. Single molecule mRNA Fluorescent In Situ Hybridization (smFISH) enables detection, counting and localization of single mRNA molecules within morphologically intact individual cells^25^. In this approach, fluorescently labeled viral and host mRNA accumulate in infected cells, whereby probe fluorescence intensity becomes a proxy for gene expression levels^26^. It therefore provides a new tool to visualize specific microbial interactions within its natural heterogeneous populations.

Here, we provide unprecedented quantification of active viral infection and host response of the cosmopolitan unicellular alga *Emiliania huxleyi* both in the laboratory and in the field. *E. huxleyi*’s massive oceanic bloom demise are considered to be terminated via infection by its large dsDNA virus EhV^3,27,28^, a member of the Nucleo-Cytoplasmic Large DNA Viruses^29^. We detected and characterized host and virus fluorescently labelled mRNA by high throughput imaging flow cytometry and tracked infection dynamics at a single virocell level. Our approach, named Virocell-FISH, provides major insights into the life cycle of lytic giant virus and its ecology, including how virocells can release virions without lysing, and lyse without releasing virions even though the host undergoes a synchronized demise.

To track the metabolic state of *E. huxleyi* during viral infection, we monitored the mRNA expression of *psbA*, a chloroplast-encoded gene of the D1 protein. D1 is a major component of the photosystem II (PSII) complex involved in the first step of the light reaction during photosynthesis. D1 has a rapid turnover, requiring constant transcription of *psbA*^30^, and is essential for optimal viral infection *E. huxleyi*^31^. Concomitantly, we followed the expression of the viral gene *mcp*, that encodes the major capsid protein expressed at the late stage of the viral program^24^ (**Fig. 1A**). Analysis of epifluorescence images of cells at 1 and 24 hours post infection (hpi) showed an increasing fraction of cells expressing viral *mcp* at 24 hpi in concert with a profound loss of *psbA* expression (**Fig. 1B, 1C**).

**Fig. 1.**
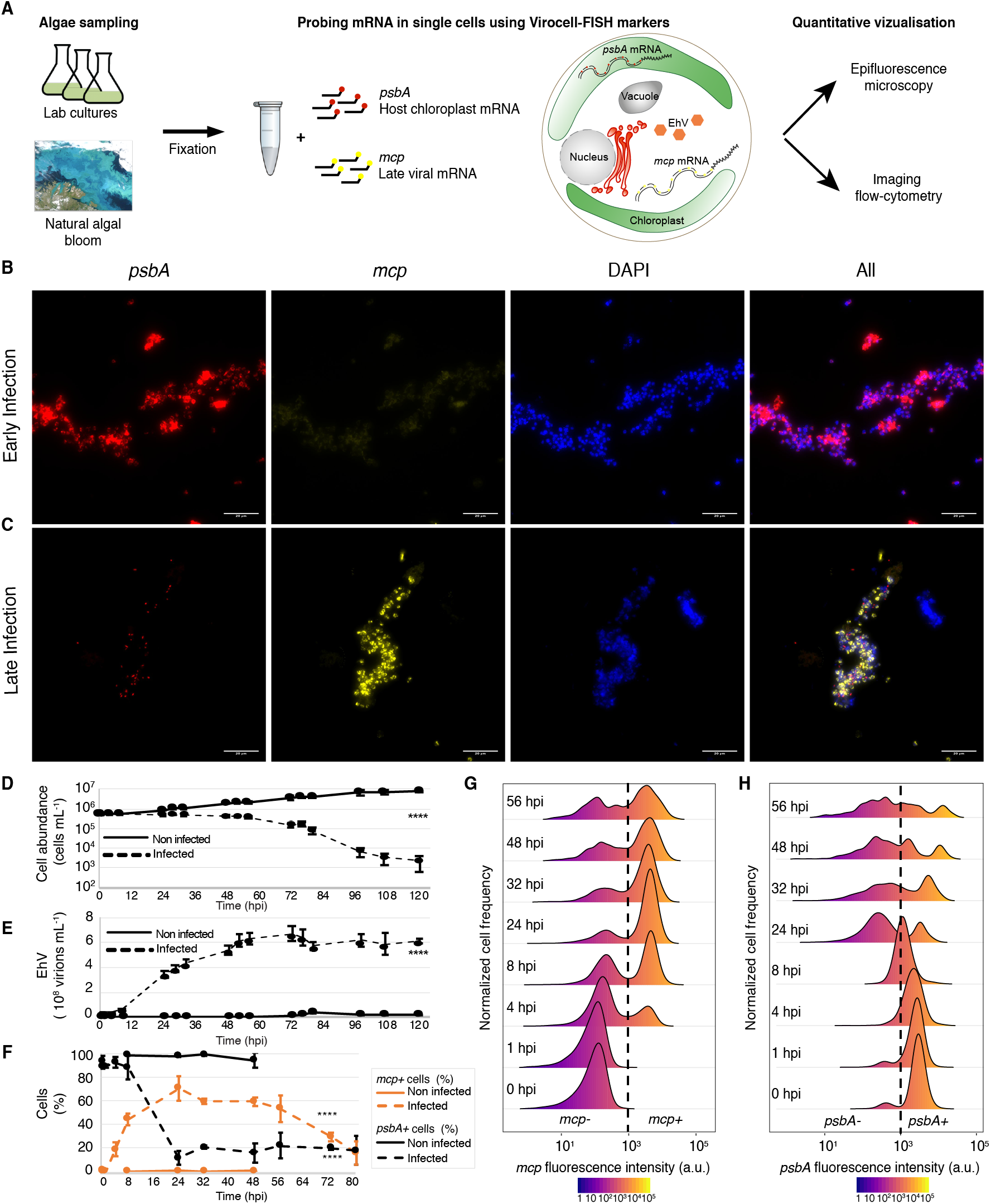
Virocell-FISH analysis enables high-throughput visualization of active viral infection in single-celled phytoplankton. (**A**) Simplified workflow of sample processing and data acquisition. After initial fixation, samples are hybridized to custom-made fluorescent probes and are subjected to either epifluorescence microscopy (high resolution) and imaging flow cytometry (high throughput) analyses, or both. (**B**, **C**) Epifluorescence images of infected *E. huxleyi* cells at 1 and 24 hpi (early and late infection, respectively) in each different channel (Scale bar = 20 μm).. (**D**, **E**) Cell and virion concentrations respectively, of infected and non-infected cultures during an *E. huxleyi* infection at high virus:host ratio. (**F**) Fraction of *mcp+* and *psbA+* cells throughout viral infection. Values are presented as the mean ± s.d., n = 3. (**G**, **H**) Distribution of *mcp* and *psbA* fluorescence intensity values across single cells of *E. huxleyi* acquired on ImageStream during a time course of viral infection. The dashed line depicts the intensity threshold used to define *mcp+* and *psbA+* cells according to the fluorescent intensity (10^3^ a.u.). **** *P* < 0.0001 tested with Linear mixed model fit by REML. T-tests use Satterthwaite's method.

In order to quantify active viral infection and host metabolic state in a high throughput manner, a highly resolved time course of *E. huxleyi* infection at high host:virus ratio was performed. Cell abundance of infected cultures remained relatively stable at 5*10^5^ cell per ml before decreasing at 48 hpi during the onset of the lytic phase (**Fig. 1D**). The first increase in extracellular concentration of virions occurred at 8 hpi before reaching a plateau at 56 hpi of 5.3*10^8^ per mL (**Fig. 1E**). Based on Virocell-FISH, samples were probed against *mcp* and *psbA,* stained with DAPI and acquired using a multispectral imaging flow-cytometer (ImageStream). *mcp*+ cells represented 19% of the population at 4 hpi, 75% at 24 hpi, plateaued at 60% between 32 and 48 hpi before declining (**Fig. 1F**). Host *psbA* expression showed that 98% of the non-infected control cells remained *psbA*+, in contrast with a sharp drop to 12% in the infected cultures between 8 and 24 hpi (**Fig. 1F**). Infected single cells, defined as *mcp*+, showed large variability in the amounts of *mcp* mRNA per cell, reflected by a wide dynamic range of intensity values in the *mcp* probe per cell (from 10^3^ to 10^5^ a.u. fluorescence), suggesting cell-cell heterogeneity in viral infection (**Fig. 1G**). Simultaneously, we detected a dual response in the population with high and low *psbA* expression (threshold at 1.1 × 10^3^ a.u. fluorescence) indicating the loss of photosynthetically active cells in a distinct subpopulation (**Fig. 1H**). Virocell-FISH therefore enables dual quantification of host and viral genes during infection dynamics across thousands of single cells.

To investigate *E. huxleyi* virocell heterogeneity during infection, we plotted host and virus mRNA expressions in parallel and revealed four distinct co-existing subpopulations (**Fig. 2A**) that we quantified through time of infection (**Fig. 2B**, **Fig. S1**). At 0 and 1 hpi, most of the cells were *mcp*−/*psbA*+ (green gate), indicating photosynthetically active cells. Control cells continuously appeared in this gate. At 4 hpi, 24% of the cells were both *mcp*+ and *psbA*+ (red gate), co-expressing viral and host genes. Two new subpopulations were clearly distinguishable at 24 hpi: 60% of the cells were *mcp*+/*psbA*− (yellow gate) as a consequence of *psbA* transcription shutdown of *mcp*+/*psbA*+ cells, and 20% of the cells were *mcp*−/*psbA*− (grey gate) and most likely originated from *mcp*−/*psbA*+ experiencing gradual photosynthetic shutdown, and cells from *mcp*+/*psbA*− that interrupted *mcp* transcription. We further compared those four subpopulations with single cell dual RNA-Seq data from^24^ and confirm that our subpopulations correspond to clear stages of viral progression as defined by kinetic classes of viral genes (**Fig. S2**). Viral release was observed within 8 hpi, yet no cell death could be detected by Sytox Green staining (**Fig. 2C)** which suggested that metabolically active infected cells (*mcp*+/*psbA*+) are responsible for early viral release. This was supported by the fact that *mcp+*/*psbA+* cells had much higher DAPI intensity than other subpopulations and showed strong positive correlation between *mcp* and DAPI intensities (**Fig. S3)**, a likely consequence of the virus-induced *de novo* nucleotide synthesis required to meet the high demand of giant viruses^32^ (**Fig. 2D**). Among *mcp*+/*psbA*− cells at 24 hpi, *mcp* and DAPI signals were strongly co-localized, suggesting overlap between viral genome dsDNA and transcripts of viral structural proteins (**Fig. S4).***psbA* intensity in infected cells (*mcp*+/*psbA*+) was higher than in non-infected cells (*mcp*−/*psbA*+) at 8 and 24 hpi (**Fig. S5)**. Higher *psbA* expression in *mcp*+/*psbA*+ could either reflect enhanced *psbA* transcription in infected cells or down regulation of *psbA* translation leading to accumulation of mRNA, as shown for plant viruses^33^ though transcript and protein levels can be uncorrelated^34^ and that at the population level cell chlorophyll intensity decreases (**Fig. S6**). Higher *psbA* in *mcp*+ population emphasized the interdependence between optimality of infection (*mcp* expression) and host metabolic regulation (*psbA* expression).

**Fig. 2.**
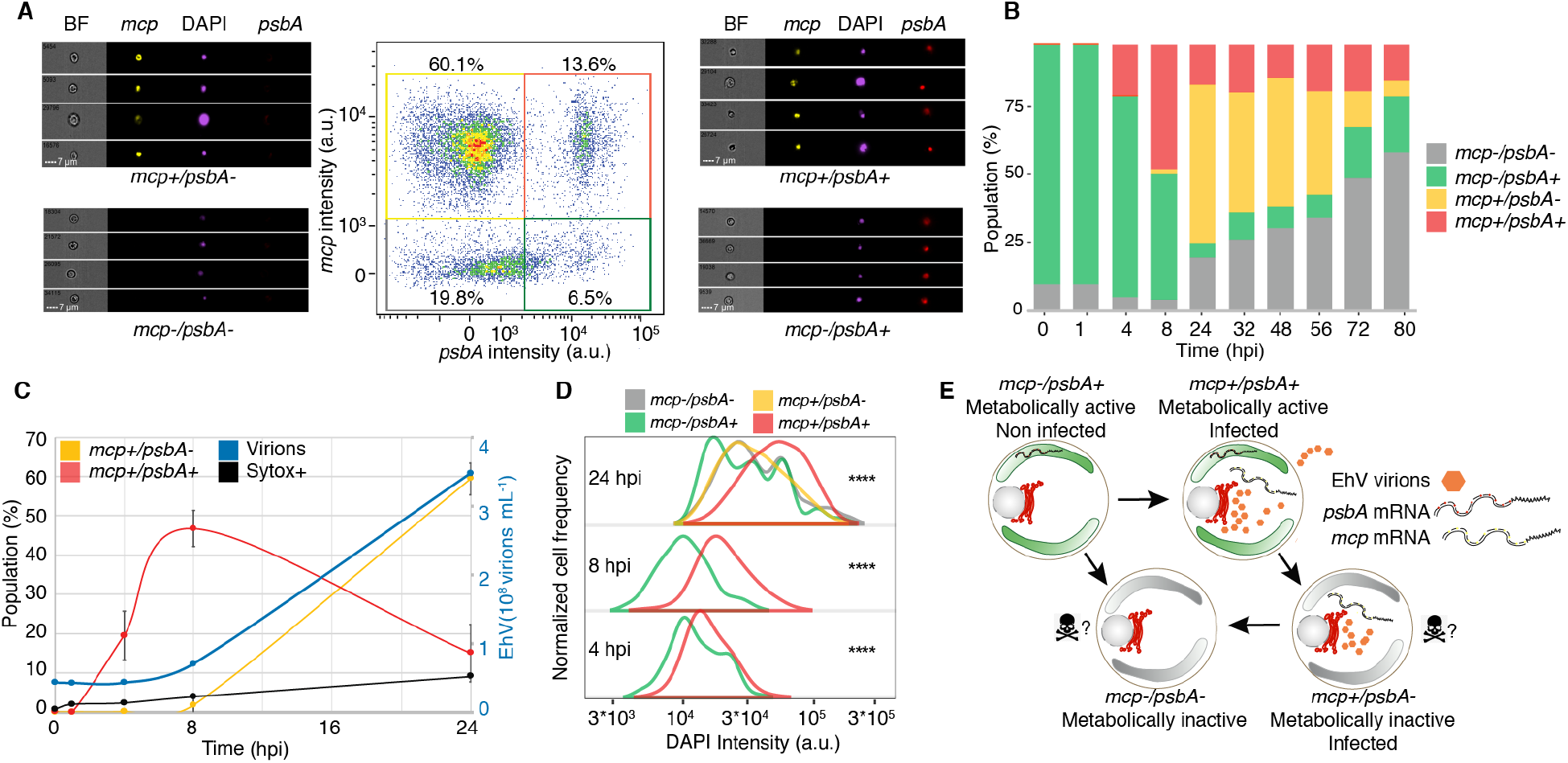
Heterogeneity in transcriptional states during viral infection reveals distinct potential cell fates. (**A**) The central panel represents co-expression of host *psbA* and viral *mcp* (X and Y axes, respectively in arbitrary units of fluorescence) in an infected *E. huxleyi* culture 24 hpi. Four subpopulations, representing distinct transcriptional states, are observed: *mcp-/psbA+* (green gate), *mcp+/psbA+* (red gate*), mcp+/psbA-* (yellow gate) and *mcp-/psbA-* (grey gate). Single-cell morphology and fluorescent signal of each subpopulation is presented, showing brightfield image (BF), *mcp* (yellow), DAPI (purple) and *psbA* (red). (**B**) Relative abundance of the four subpopulations throughout the course of infection. (**C**) Close-up on virion production (virions ml^−1^), cell death (Sytox+), and *mcp*+ dynamics in the 24 first hours post infection. (**D**) Comparison of DNA content based on DAPI intensity between the four subpopulations (with more than 25 events per population) throughout infection. Comparison of DAPI intensity was with a Linear mixed model fit by REML. T-tests use Satterthwaite's method. * *P* < 0.05, **** *P* < 0.0001. (**E**) Schematic model of potential cell fates of the different subpopulations.

Interestingly, based on our intracellular virocell infection dynamics, we quantify fundamental parameters in the life cycle of a marine giant viruses. We estimate that the eclipse phase, in which viruses have penetrated the host but no virions are produced^35^, occurs within the first 4 hours, based on the induction of *mcp*+/*psbA*+ cells, without virion increase (**Fig. 2C**). The onset of the maturation phase, during which viral progeny is released, occurs within 8 hours. The distinct infection states can be associated with unique cell fates: metabolically active infected cells (*mcp+/psbA+*) are responsible for early viral release most likely through budding whilst *mcp*+/*psbA*− likely contribute less to this process. The *mcp*−/*psbA*− subpopulation, correlated with Sytox Green staining (**Fig. S1B**), suggests that a substantial fraction of the cells lyse without releasing viruses after 56 hpi. These cells can die from accumulation of DNA damage^36^, cytotoxic viral glycosphingolipids coming from neighboring infected cells^37^, or by abortive infection. The accumulation of these cells during lytic phase can originate from either the *mcp*+/*psbA*+ subpopulation that halts *mcp* transcription before lysis, or from the *mcp*−/*psbA*+ subpopulation that shuts down *psbA* transcription^24^. At later time point, the *mcp*−/*psbA*+ cells might serve as the seed for the resistant phenotypes often observed in small subpopulation that recovered from viral infection^38^. The transition between different phenotypic states could be mediated by an array of infochemicals, such as viral glycosphingolipids^37,39^, or by extracellular vesicles^40^ (**Fig. 2E)**. Virion egress from the host cells without induction of cell death may provide an optimal co-existence mechanism during bloom initiation phase^41^.

The consequences of viral infection in the ocean are not limited to the cell fate of the individual host cell. Viral infection can also lead to aggregate formation, mediated by production of transparent exopolymer particles (TEP) consisting of acidic polysaccharides, which are classically quantified in bulk by staining with Alcian Blue^42^. Aggregates accelerate sinking of biomass, contribute to “marine snow” and enhance carbon export to the deep sea^3^. In order to assess the contribution of viral infection to aggregate formation, we quantified aggregated *E. huxleyi* cells and characterized the infection state among individual cells within the aggregates using Virocell-FISH (**Fig. 3A**). Upon infection, up to 40% of the algal cells (DAPI positive fraction) were found within aggregates (**Fig. 3B**). At the early stage of infection, aggregates are rare (<10%), mostly infected (75% of the aggregates were *mcp*+) and photosynthetically active (100% were *psbA*+) (**Fig. 3D, 3E**). At the onset of the lytic phase (48 hpi), aggregates were highly abundant, less infected (25% were *mcp*+) and less photosynthetically active (30% were *psbA*+). In infected cultures at 8 hpi, the fraction of *mcp*+ aggregates was higher than the fraction of *mcp+* single cells (75% and 45%, respectively, **Fig. 1F and Fig. 3D**). We hypothesize that aggregates enriched with infected cells may favor sinking of newly produced virions out of the photic zone and prevent further dissemination of viruses in the bloom, suggesting a host defense strategy^3,28^. The overall presence of infected cells in aggregates corroborates previous observations of TEP-attached viruses^42,43^. This could further explain the estimates of 10^9^ viruses per gram of marine sediment, some being infectious, hence may serve as a possible inoculum to infect subsequent blooms^44,45^.

**Fig. 3.**
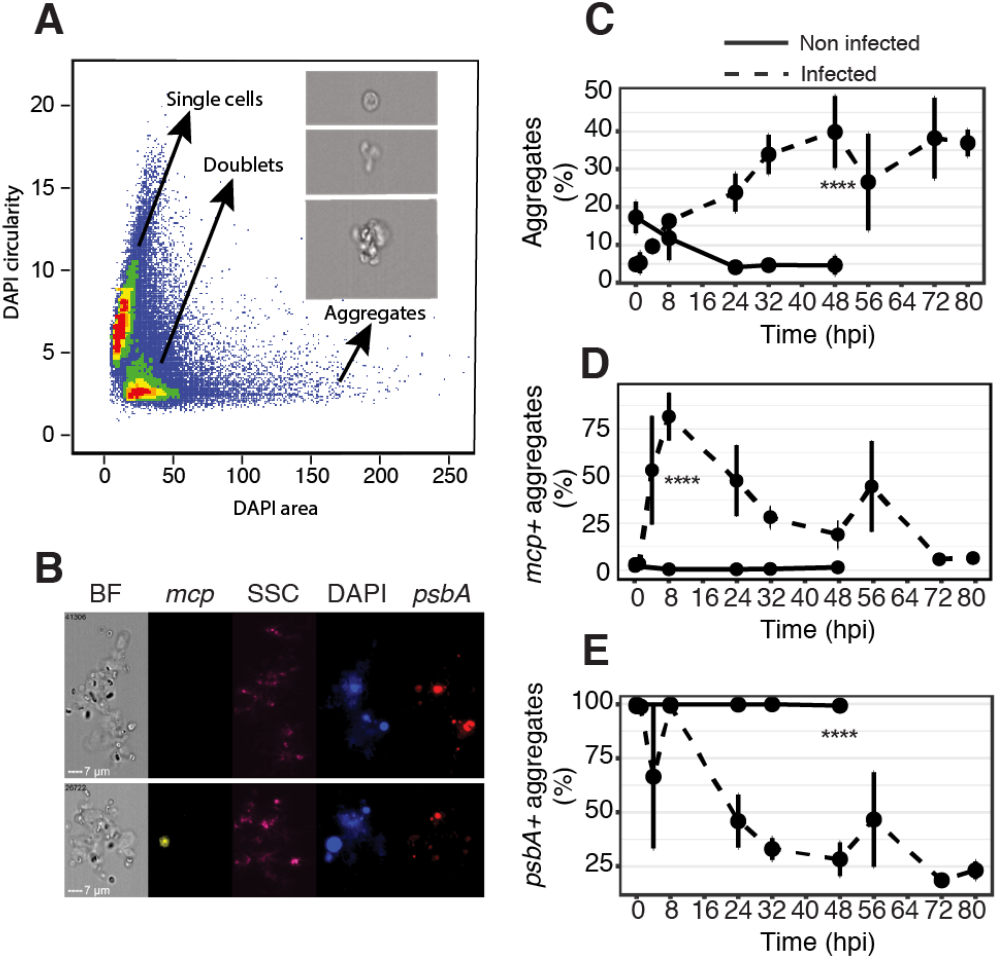
Quantification of viral infection within aggregates formed during infection dynamics. (**A**) Based on the DAPI mask area and circularity (the degree of the mask’s deviation from a circle), three populations are identified as single cells, doublets and aggregates (aggregates are defined by a DAPI area above 60 μm^2^). (**B**) Imaging flow-cytometry images of non-infected (top) and infected (bottom) aggregates in the brightfield (BF), *mcp* (yellow), Side Scatter (SSC as a proxy for calcification, pink), DAPI (blue), and *psbA* (red) channels. (**C**) Fraction of aggregates (DAPI area above 60 μm^2^) out of all DAPI+ events in infected and non-infected cultures. (**D**, **E**). Quantification of *mcp+* and *psbA+* aggregates respectively, through time across three replicates between infected and non-infected cultures. **** *P* < 0.0001 (t-test).

One of the fundamental knowledge gaps in aquatic virology is the ability to quantify active viral infection in natural populations and to mechanistically resolve the viral impact on microbial ecosystems. We therefore conducted a mesocosm experiment in a Fjord near Bergen, Norway, and collected samples for Virocell-FISH analysis across different phases of an *E. huxleyi* bloom. We quantified viral infection specifically in *E. huxleyi* cells by using a 28S ribosomal probe unique to that algae, a *psbA* probe as a proxy for host metabolic state and a probe targeting viral *mcp* transcripts (**Fig. 4A**). Virocell-FISH analysis conducted by imaging flow cytometry successfully revealed the fraction of infected *E. huxleyi* cells during bloom succession, based on over 5000 cells at each time point (**Fig. S7**). From the initiation of the bloom to the peak in *E. huxleyi* abundance, infection level (% of *mcp*+ cells) was minimal (<1%). A tipping point was detected in which the fraction of infection cells rose from 1.51% on day 17 to 23.22% on day 18 while cell abundance dropped at the onset of bloom demise (**Fig. 4B**). Strikingly, the maximal fraction of infected cells on the demise phase, did not reach beyond 27% of *E. huxleyi* population even at the highest levels of *E. huxleyi* infection. Furthermore, *mcp*+ cells displayed higher DAPI intensity than *mcp*− cells (**Fig. 3C**), confirming laboratory results regarding the induction of *de-novo* nucleotide synthesis in infected cells. Quantification of co-expression of *mcp* and *psbA* over the course of the bloom (**Fig. S8**) revealed a major drop in *mcp*−/*psbA*+ cells between days 16 and 18, accompanied by an increase in *mcp*+/*psbA*- cells. From day 19 onwards a large fraction of the *E. huxleyi* population was *mcp*−/*psbA*- resembling increase in the fraction of cell death during bloom demise (**Fig. 4F**). These results suggest algal bloom demise can be synchronized despite co-existence of diverse virocell states and only a subpopulation of infected cells. We suggest that other mortality agents might act in concert with viral lysis to control synchronized bloom demise at a later stage. These top-down regulators may include pathogenic bacteria^46^, eukaryotic parasites and grazers^3,47,48^, programmed cell death of non-infected bystander cells mediated by release of viral-derived infochemicals^37,39^ or extracellular vesicles^40^. Alternatively, a highly synchronized viral infection and release might have occurred in a large fraction of the *E. huxleyi* population in a narrow time frame between day 17 and 18. Indeed, recent reports demonstrated that viruses are intrinsically synchronized with the daily rhythms of their photosynthetic host both in controlled lab experiments and in natural populations^49,50^.

**Fig. 4.**
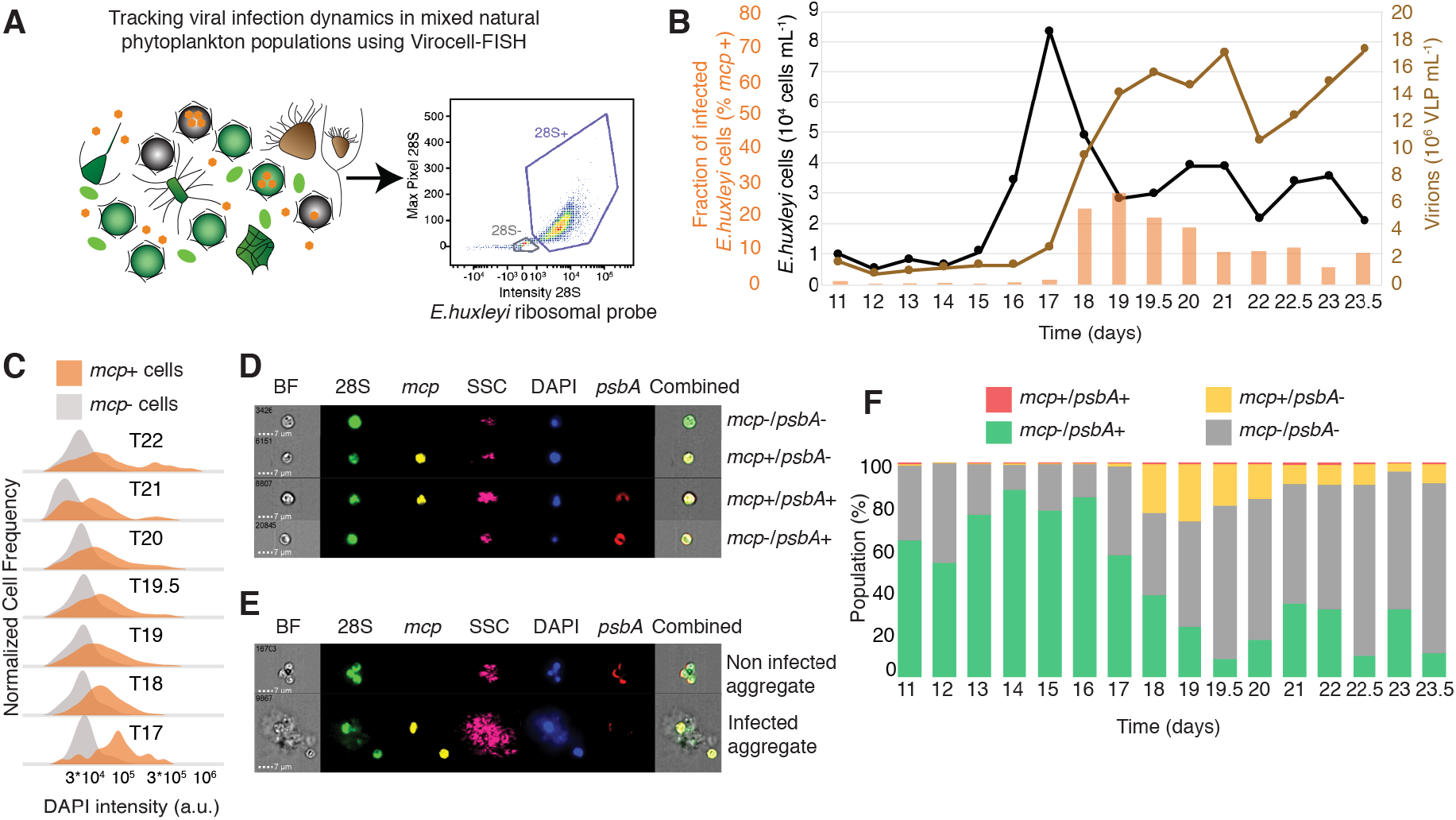
Visualization of active viral infection during a natural *E. huxleyi* bloom. (**A**) Applying Virocell-FISH to track viral infection of *E. huxleyi* cells within mixed natural microbial populations. Viral mRNA was targeted using the *mcp* probe whilst host mRNA was monitored using *psbA*; an additional probe specifically targeting the 28S ribosomal RNA of *E. huxleyi* was used to identify our host of interest, based on the intensity and max pixel of the probe signal. (**B**) Fraction of infected (*mcp*+) *E. huxleyi* cells (orange), abundance of calcified *E. huxleyi* cells (black) and EhV like particles (brown) throughout bloom succession (days post inoculation) in a mesocosm experiment. Active viral infection in *E. huxleyi* cells was measured using Virocell-FISH; *E. huxleyi* and EhV-viral like particles abundance were measured by flow-cytometry. Days with 0.5 digit represent evening samples. (**C**) Comparison of DAPI intensities between infected and non-infected *E. huxleyi* populations throughout bloom succession. (**D**, **E**) Visualizing subpopulation of *E. huxleyi* single cells and aggregates respectively, using ImageStream. (**F**) Quantification of the relative abundance of the four subpopulations in single cells throughout the bloom succession, by co-probing of host and viral mRNA (*psbA* and *mcp* respectively).

Our approach opens up new avenues to study individual virocells in the marine environment, by following infection dynamics both intra- and extracellularly of morphologically intact cells. It unravels fundamental aspects of the life cycle of a large virus by linking scales between single cell and population level dynamics, showing viral increase without cell lysis, and cell lysis without viral increase. Such high-resolution examination of host and virus transcriptional states reveals cell-to-cell heterogeneity in viral infection, mediated by the dependence on the host metabolic state, despite synchronized bloom demise. Concomitantly, it can inform about viral hijacking mechanisms of the host photosynthetic machinery. Coupling enhanced viral and photosynthetic transcriptions may require to maintain photosynthetic activity to energize the redirection of carbon flux from the Calvin cycle to the pentose phosphate pathway in order to enhance production of NADPH required for *de novo* DNA synthesis^51^ to meet the high demand of DNA synthesis of giant viruses and to enhance the antioxidant capacity under oxidative stress^52^. Therefore, we propose that a functional convergence arises among viruses infecting phototrophs from diverse evolutionary origin, whereby cyanophages^53^, small RNA plant viruses^54^ and large algal dsDNA viruses share the strategy to manipulate their photosynthetic host metabolism by different mechanisms. Quantifying the virocells infection and metabolic states, through their diverse phenotypes will help integrate viral-dependent processes in ecosystem models by providing necessary numbers of infection dynamics, such as the fraction of actively infected cells, resistant cells, the nature of sinking biomass, as well as their genetic composition as a function of time and other stresses^55^. Finally, this quantitative approach can be expanded to diverse host-pathogen and host-symbiont systems with important ecological significance, adding a mechanistic dimension to the field of marine microbial ecology.

## Acknowledgments

We thank S. Itzkovitz, L. Farack, S. Ben-Moshe for guidance in smFISH protocols, analysis, and fruitful discussions; D. Hirsch for training on the epifluorescence microscope. The AQUACOSM-VIMS consortium for the mesocosm experiment and all Vardi lab members for discussion. We particularly thank G. Schleyer and D. Schatz for critical reading, and R. Avraham for useful critical feedback

## Author contributions

F.V. and A.V. designed and conceptualized the study. U.S. developed smFISH on *E. huxleyi* and F.V. implemented it in high throughput. Z.P. helped with ImageStream analysis F.V. prepared the cell cultures, performed all acquisition and data analysis. F.V. prepared the figures and wrote the manuscript with A.V. and input from U.S.; A.V. supervised the study

## Data and materials availability

All data, code, and materials used in the analysis are available upon demand.

## Supplementary Materials

Materials and Methods

Table S1

**Table S1. Calculation of virus:host ratio using the MPN method**

Figures S1-S13

**Fig. S1. Dynamics of subpopulations during infection in EhV201**

**Fig. S2. Comparison of Virocell-FISH with single-cell RNA-Seq**

**Fig. S3.** *mcp* **and DAPI intensities in single cells throughout infection**

**Fig. S4. Morphology of metabolically active infected and non-infected cells**

**Fig. S5. Temporal dynamics of** *psbA* **intensities in** *psbA* **+ subpopulations**

**Fig. S6. Chlorophyll mean intensity during infection**

**Fig. S7. Adapting Virocell-FISH to target specifically** *E.huxleyi* **in environmental samples**

**Fig. S8. Defining** *mcp*/*psbA* **subpopulations in environmental samples.**

**Fig. S9. Pipeline of single cell identification in ISX (Only SM)**

**Fig. S10. Comparison of the Virocell-FISH method with or without RNAase treatment (Only SM)**

**Fig. S11. Quantifying infected cells in microscopy data using CellProfiler (Only SM)**

**Fig. S12. Gene expression of** *psbA* **and** *mcp* **in single cell RNA-seq sequencing data (Only SM)**

**Fig. S13. Flow cytometry gates used for** *E. huxleyi* **cells in the field(Only SM)**

