## Supplementary Material for "Visualizing active viral infection reveals diverse cell fates in synchronized algal bloom demise"

**This PDF file includes:**

Materials and Methods

Figs. S1 to S13

Table S1

### Materials and Methods

Culture growth and viral infection

The non-calcifying *E. huxleyi* strain CCMP 2090 was used for this study. Cells were cultured in K/2 medium with antibiotics (Ampicillin and Kanamycin) and incubated at 18°C with a 16:8h light-dark illumination cycle. A light intensity of 100 μmol photons $m^{-2}$ $s^{-1}$ was provided by cool white light-emitting diode lights. All experiments started with exponential phase cultures (5*10^5 cells $mL^{-1}$). The virus used for this study is EhV201 propagated on CCMP2090 with antibiotics. Five day before infection, Most Probable Number assays were performed to assess the fraction of infectious viruses in the stock (*48*). *E. huxleyi* was infected with 5:1 multiplicity of infection (MOI) ratio of infectious virus per cell, hence guaranteeing that all cells encountered an infectious particle at 1 hour post infection (hpi). Time course infections were done on the same day, in triplicates. A second MPN experiment was performed on the day of the experiment with the same virus and same algae to assess the exact MOI at the beginning of the experiment. (**Table 1**).

Enumeration of algal cell abundance, cell death and viral abundance

Cells were monitored and quantified using an Eclipse (iCyt) flow cytometer. Cells were identified by plotting the chlorophyll fluorescence (excitation (ex): 488nm and emission (em): 663-737nm) versus side scatter. For extracellular viral counts, samples were stained with SYBR gold (Invitrogen) that was diluted 1:10000 in Tris-EDTA buffer, incubated for 20 min at 80°C and cooled to room temperature (RT). Samples were analyzed by an Eclipse flow cytometer (ex: 488 nm and em: 500-550 nm), a minimum of 50,000 events was collected. To analyze cell death, samples were stained with a final concentration of 1 μM Sytox Green (Invitrogen), incubated in the dark for 1h at RT and analyzed by an Eclipse flow cytometer (ex : 488 nm and em: 500-550nm).

An unstained sample was used as a control to eliminate the background signal. The data was exported and analyzed in R.

*mcp*, and *psbA* probe design and conjugation for Virocell-FISH

Fasta sequences of the target genes were first submitted to Stellaris Probe Designer to obtain potential probes. We designed 48 probes per gene with a probe length of 20 nucleotides. Probes with more than 70% GC content were discarded. To discard off-targets and decrease nonspecific signal, each probe sequence was blasted against the *E. huxleyi* transcriptome and EhV201 genome. Probes with an off-target gene matching over 17 nucleotide were discarded. Validated probes were ordered with 3' amine groups through the Custom Oligo Service of BioSearch Technologies (see **Probe_sequence.xlsx**). Fluorophores with succinimidyl ester group were ordered from Click Chemistry Tools: Tetramethylrhodamine (TMR), Alexa594 (AF594) and sulfo-Cyanine5 (Cy5). Probes were coupled to fluorophores and purified according to[1].

Sample fixation and hybridization for Virocell-FISH in laboratory samples

At each time point, 50 mL of each flask were fixed in cold 1% paraformaldehyde final concentration, and incubated for 1h at 4°C with gentle agitation. Each sample was then centrifuged for 2 min at 4°C and 3000g. The supernatant was discarded, the pellet resuspended in 1 mL of cryopreservant solution (prepared in 1X PBS containing 4% paraformaldehyde and 30% sucrose), transferred to a 1.7 mL Eppendorf and incubated 1h at 4°C with agitation. Tubes were then centrifuged for 2 min at 4°C and 3000g, the supernatant removed, and stored at 80°C until hybridization.

A day before ImageStreamX acquisition, selected samples were thawed and chlorophyll extracted by a first wash using 900 µL of 70% ethanol applied for 3 min, followed by centrifugation 3 min at 3000g and removal of supernatant. A second wash was performed with 900 µL of 100% ethanol applied for 3 min, followed by centrifugation of 3 min at 3000g and removal of supernatant. Samples were treated with 500 µL ProteinaseK at 10 µg/mL final concentration (Ambion #AM2546) for 10 min and washed with 3 minutes at 3000g centrifugation. Samples were then resuspended in 50 µL of hybridization buffer (17.5% formamide concentration) containing equal concentrations of the different target genes at 0.1ng/mL final concentration. *Mcp* probe was conjugated to TMR, and *psbA* was coupled to Cy5. Hybridization was performed overnight in 30°C shakers.

The day of ImageStreamX acquisition, hybridization buffer was washed away by 3 min centrifugation at 3000g. Samples were stained for dsDNA with 500 µL of GLOX buffer (prepared in nuclease free water with 0.4% final concentration of glucose, 2X final concentration SSC Ambion #AM9765, and 10 mM final concentration of Tris pH 8.0) with DAPI in 10µg/mL final concentration (except the single stains). DAPI staining was done for 30 min in 30°C, followed by centrifugation 3min at 3000g, removal of supernatant and resuspension in 40µL of GLOX buffer before being acquired in the ImageStreamX.

The following day, samples were resuspended in 300 µL of GLOX with the 3,3'-Dihexyloxacarbocyanine Iodide membrane stain at final concentration of 1µM (ThermoFisher Catalog number D273). Excessive dye was removed by 3 min centrifugation at 3000g and removal of supernantant, followed by resuspension in 4 µL GLOX. 2 µL were deposited on a microscope slide with 2 µL of antibleach solution, and the sample was imaged with the epifluorescent microscope.

Imaging Flow Cytometry acquisition, compensation and analysis for laboratory samples

Samples were acquired using the ImageStreamX MarkII machine (ISX, Amnis, Luminex). Three excitation wavelengths were used: 405 nm (DAPI - Channel 7- 50 mW), 561 nm (TMR - Channel 3 – 200 mW) and 642 nm (Cy5 - Channel 11 – 120 mW). For each sample, at least 50,000 cells were acquired and imaged with 60X magnification. Single stained samples were acquired using the compensation wizard of ISX. On average, less than 10 µL of 2x10$^{\wedge 6}$ cell stock concentration were necessary to collect the optimal amount of cells, except towards the end of the experiment where cells were scarce.

Data was analyzed using IDEAS6.2 (Amnis, Luminex). The compensation matrix was built using the IDEAS wizard and manually checked, before being applied to all the acquired files. Based on the Area (the number of microns squared in a mask) and Circularity (the degree of the mask's deviation from a circle) of DAPI three populations are identified as single cells (mainly DAPI area < 60 a.u), doublets and aggregates (mainly DAPI area > 60 a.u). Single cells were additionally selected in the same focal plane using the BF gradient and contrast (both gradient and contrast measure the sharpness quality of an image by detecting large changes of pixel values in the image). All gates were defined on a single file before being applied the total data set (See **Fig. S9**). Each file was then manually inspected to check the accuracy of single cell and aggregates gating. All the data (fluorescent intensities, morphological features, populations) was then exported for each cell of each file for analysis in R.

RNAse treatment

*E.huxleyi* cultures were infected with EhV201 at low MOI and fixed in 1% paraformaldehyde for Virocell-FISH 24 hours post infection. Two samples were used to compare *mcp* signal with and without RNAase treatment. The sample with RNAse treatment was treated with RNAse before the Proteinase-K treatment, with RNAse from Thermo-Fisher (cat. EN0531) at a final concentration of 10 μg/mL in a reaction volume of 200 μL of infected *E.huxleyi* culture, for 20 min at 37 degrees. Both samples were then processed as usual for *mcp* staining and acquired in the ImageStream. Application of RNAase treatment on infected samples shows heavily reduced the *mcp*$_{+}$ signal confirming that the probes bind to RNA and not DNA (**Fig. S10**)

Epifluorescence microscope acquisition and analysis

Slides were acquired using a Nikon inverted fluorescence microscope Eclipse Ti2 Series and imaged with a Ixon Ultra 888 camera with 100X magnification, using the Nikon NIS Element Advanced Research Software. Illumination time for each excitation was adjusted at the beginning of each batch acquisition and not modified after that. Each image is composed of 15 0.3 μm stacks. Images were exported in .nd2 format and inspected in Fiji 2[2]. Stacking was performed for each file (Image>Stacks>Z-Project>MaxIntensity) on maximum 10 stacks. Mean intensity in the background was measured for each channel (Analyze>Measure) and subtracted (Process>Math>Subtract).

To estimate fraction of infected cells we used CellProfiler[3] using the PercentPositive pipeline, separately for each probe after doing a max projection and background subtraction for each channel separately (See **Fig. S11)**.

Single Cell Transcriptomics Analysis

Based on single cell dual transcriptomics during *E. huxleyi* infection by *EhV201*[4], we plotted the abundance, relative expression with respect to the total RNA pool, and frequency of *psbA* versus *mcp* which we probed in the smFISH probes analysis (**Fig. S12**). Furthermore, we were able to map the single cell transcriptomics infection states (*metacell*) into each of the Virocell-FISH transcriptional states that were based on *psbA* versus *mcp* co-expression on a single cell resolution. (**Fig. S2**).

Mesocosm experimental setup

The mesocosm experiment – Aquacosm-viral induced microbial succession (Aquacosm-VIMS) – was carried out over 24 days (May 24 – June 17, 2018) in Raunefjorden at the University of Bergen's Marine Biological Station Espegrend, Norway (60.27° N; 5.22° E). The experiment consisted of seven enclosure bags made of transparent polyethylene (11 $m^3$, 4 m deep and 2 m wide, 90% photosynthetically active radiation) mounted on floating frames and moored to a raft in the middle of the bay. Each bag was filled with surrounding fjord water and supplemented with nutrients at days 0-7 and 13-17 at a nitrogen:phosphorous ratio of 16:1 (1.6 µM $NaNO_3$, 0.1 µM $KH_2PO_4$. In days 6, 7 and 13 only $NaNO_3$ was added). The water in each bag was continuously mixed by pumping air to the bottom of each bag. Samples for flow cytometric counts were taken twice a day, morning (7 am) and evening (8-9 pm) using 50 mL centrifugal tubes and following filtration using a 40 µm cell strainer. Calcified *E. huxleyi* were identified using the Eclipse flow-cytometer based on high side scattering and high chlorophyll content (**Fig. S13**).

Virocell-FISH specificities for field samples

Sample fixation, storage and initial chlorophyll washes of field samples was identical to laboratory samples. In order to identify *E. huxleyi* cells, we designed a single probe specific to the 28S region of *E. huxleyi*[5] (EG28-03, 5′-TAAAGCCCCGCTCCCGGGTT-3′, bound to C3-Fluorescein (ex/em = 490/525nm). Two helper probes were used (Helper A, 5'-GCCAGGACGGGAGCTGGCCG-3' and Helper B, 5'-GAGGCGCGGCGCCGAGGCGC-3'). 28S, HelperA, HelperB were used at final concentration of 0.5µM, at the same time as the *mcp* and *psbA* probes at 0.1ng/mL final concentration. All samples were stained in 50µL of 40% formamide hybridization buffer, incubated at 37 degrees overnight. Samples were acquired using the ImageStreamX MarkII machine (ISX, Amnis, Luminex). Four excitation wavelengths were used: 405 nm (DAPI - Channel 7- 50 mW), 488 nm (AF488 - Channel 2 - 200mW),561 nm (TMR - Channel 3 – 200 mW) and 642 nm (Cy5 - Channel 11 – 120 mW).

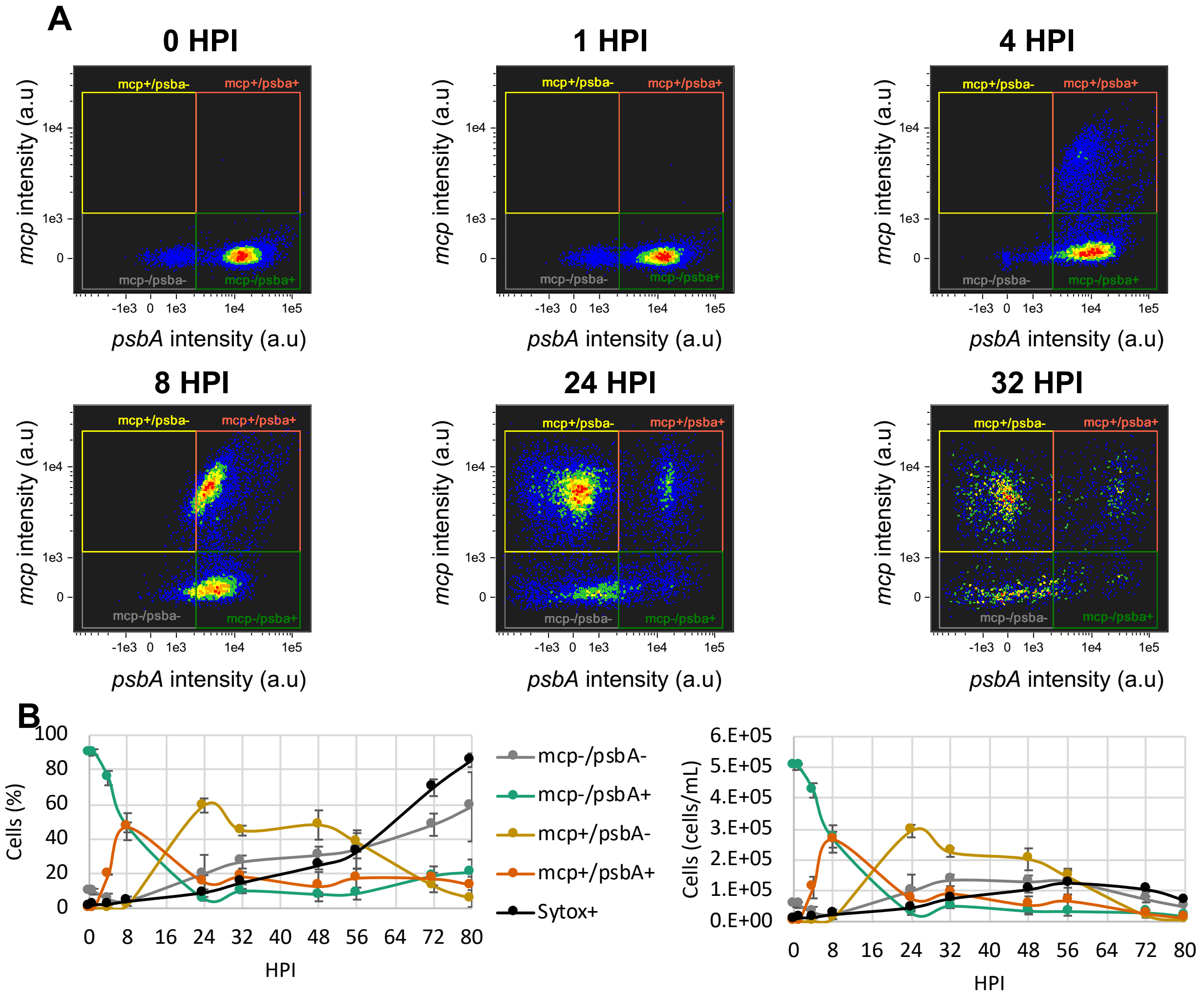


**Fig. S1. Dynamics of subpopulations during infection in EhV201**. (**A**) To investigate *E. huxleyi* virocell heterogeneity during infection, we plotted host and virus mRNA expressions in parallel at different time points of hours post infection (hpi). The X axis represents the value of the probe intensity (in fluorescent arbitrary units) targeting the *psbA* host gene, and the Y axis represents the value of the probe intensity targeting the viral *mcp* gene. Using a threshold of approximately $10^{\wedge 3}$ arbitrary units to define *mcp*+ and *psbA*+ cells, we define four subpopulations as a combination of *mcp* and *psbA* signals. (**B**) Proportion of cells in each gate through time (left panel) and absolute abundance in cell/mL of cells in each gate (right panel), including Sytox+ dynamics.

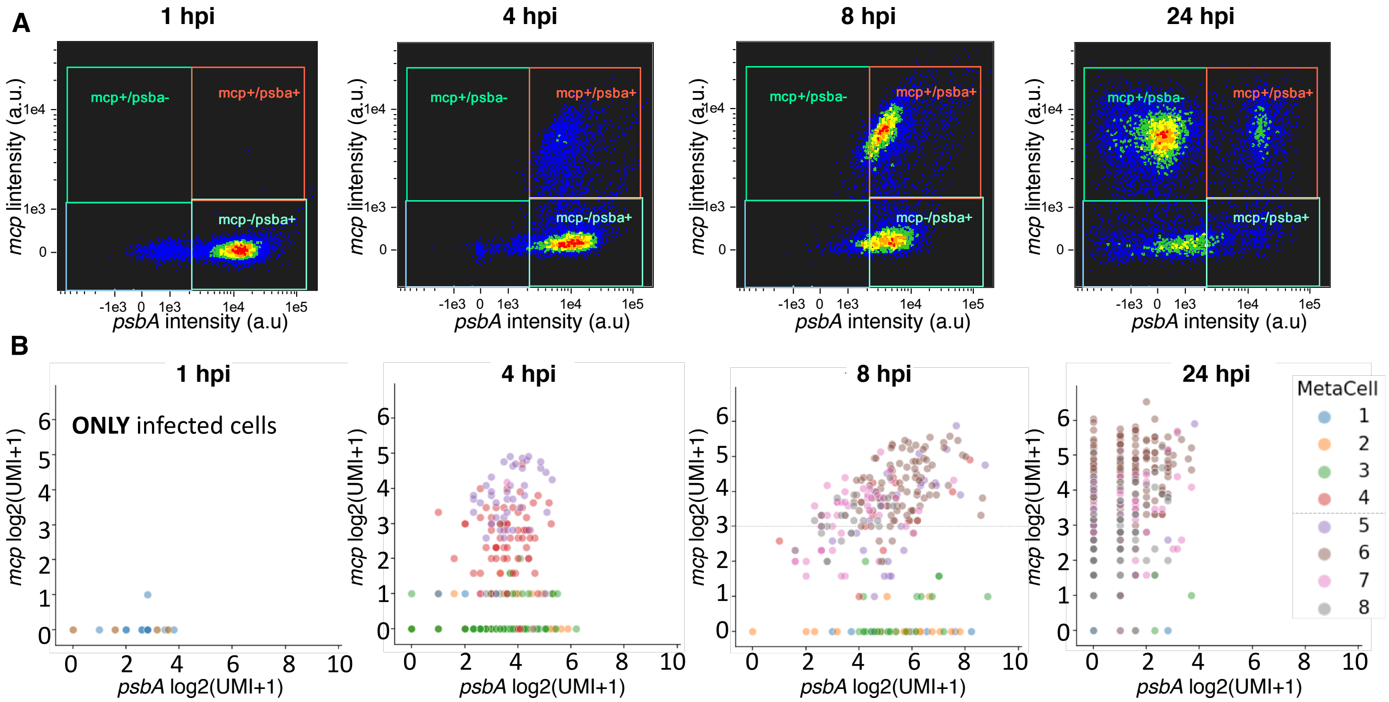


**Fig. S2. Comparison of Virocell-FISH with single-cell RNA-Seq.** (**A**) Time course of transcriptional states in EhV201 based on imaging flow cytometry and Virocell-FISH with *psbA* (x axis) and *mcp* (y axis) probes. (**B**) Time course of single cell RNA-Seq for the same time points of an EhV201 infection[4]. Each dot is an infected cell and is plotted with respect to the number of unique *mcp* and *psbA* mRNA counts (Unique Molecular Identifier, or UMI). Each cell is colored by the metacell it belongs to, where a metacell can be defined as a group of infected cells that form a cohesive group based on their total individual RNA expression. Metacell 1 (blue) represents cells in early stage of the viral program, and metacell 8 (grey) represents cells in the last stage of the viral program. For example at 4 hpi, most of the cells in the *mcp-*/*psbA+* gate belong to metacells 1-3, whereas cells in the *mcp+*/*psbA+* gate correspond to cells belonging to metacells 4-7.

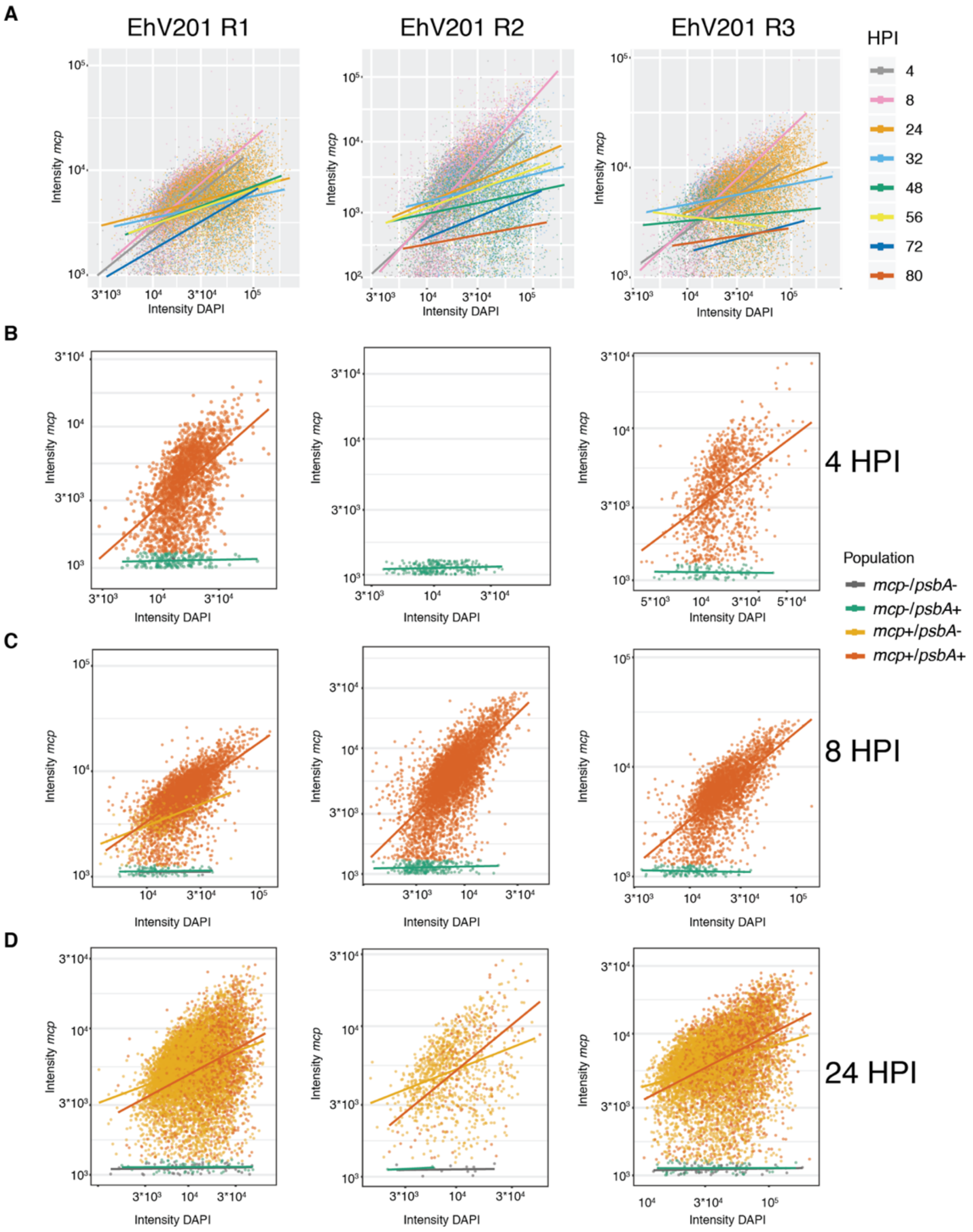


**Fig. S3. *mcp* and DAPI intensities in single cells throughout infection.** (**A**) DAPI intensity versus *mcp* intensity of single cells in *EhV201* Replicate 1,2,3. Colors represent different timepoints. Slopes were plotted based on the smoothed "lm" function (geom_smooth() in R). (**B**) DAPI versus *mcp* for *EhV201* Replicate 1,2,3 at 4 hpi colored per transcriptional subpopulation. (**C**) DAPI versus *mcp* for *EhV201* Replicate 1,2,3 at 8 hpi colored per transcriptional subpopulation. (**D**) DAPI versus *mcp* for *EhV201* Replicate 1,2,3 at 24 hpi colored per transcriptional subpopulation.

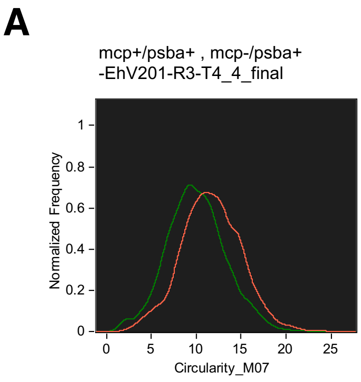


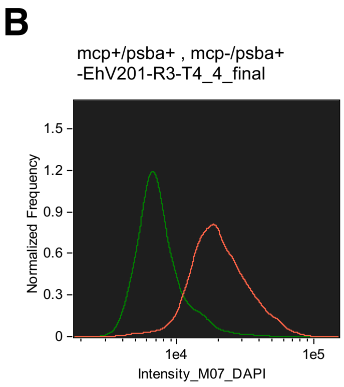


Circularity_M07

| Population | Count | %Gated | Mean |
|---|---|---|---|
| mcp+/psba+ & SingleCellFoc... | 3280 | 100 | 11.66 |
| mcp-/psba+ & SingleCellFoc... | 4399 | 100 | 9.998 |

Intensity_M07_DAPI

| Population | Count | %Gated | Mean |
|---|---|---|---|
| mcp+/psba+ & SingleCellFoc... | 3280 | 100 | 22861.72 |
| mcp-/psba+ & SingleCellFoc... | 4399 | 100 | 8069.53 |

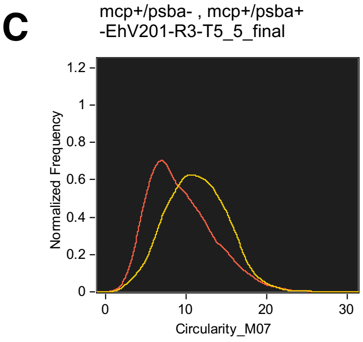


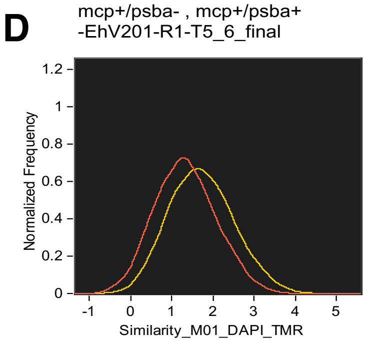


Circularity_M07

| Population | Count | %Gated | Mean |
|---|---|---|---|
| mcp+/psba- & SingleCellFoc... | 6532 | 100 | 11.31 |
| mcp+/psba+ & SingleCellFoc... | 2374 | 100 | 9.399 |

Similarity_M01_DAPI_TMR

| Population | Count | %Gated | Mean | Median | Std. Dev. |
|---|---|---|---|---|---|
| mcp+/psba+ & SingleCellFoc... | 2017 | 100 | 1.334 | 1.316 | 0.7352 |
| mcp+/psba- & SingleCellFoc... | 7802 | 100 | 1.725 | 1.694 | 0.7777 |

**Fig. S4. Morphology of metabolically active infected and non-infected cells.** (**A**, **B**) Analysis of DAPI circularity and intensity respectively in *mcp+/psbA+* cells (red) and *mcp-/psbA+* (green) 8 hpi in biological replicate 3. (**C**, **D**). Analysis of DAPI circularity and colocalization between DAPI and *mcp* signals respectively in *mcp+/psba-* cells (yellow) and *mcp+/psbA+* (orange) 24 hpi in biological replicate 3.

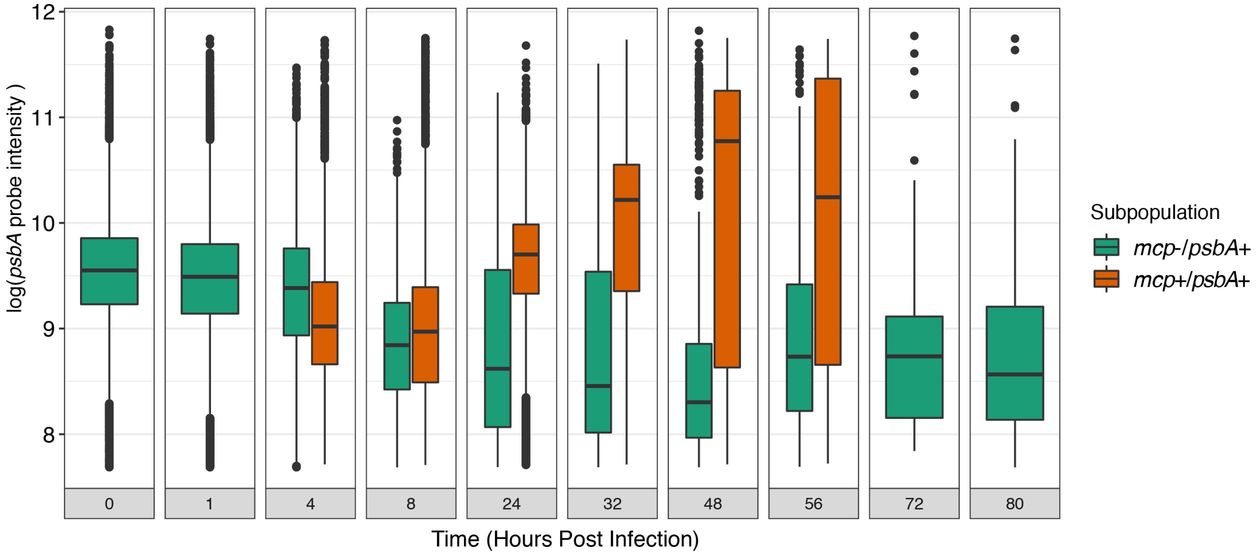


**Fig. S5. Temporal dynamics of *psbA* intensities in *psbA*+ subpopulations.** Box plot of *psbA* probe intensities in *mcp*+/*psbA*+ (orange) and *mcp*-/*psbA*+ (green) sub population over time of infection (hpi).

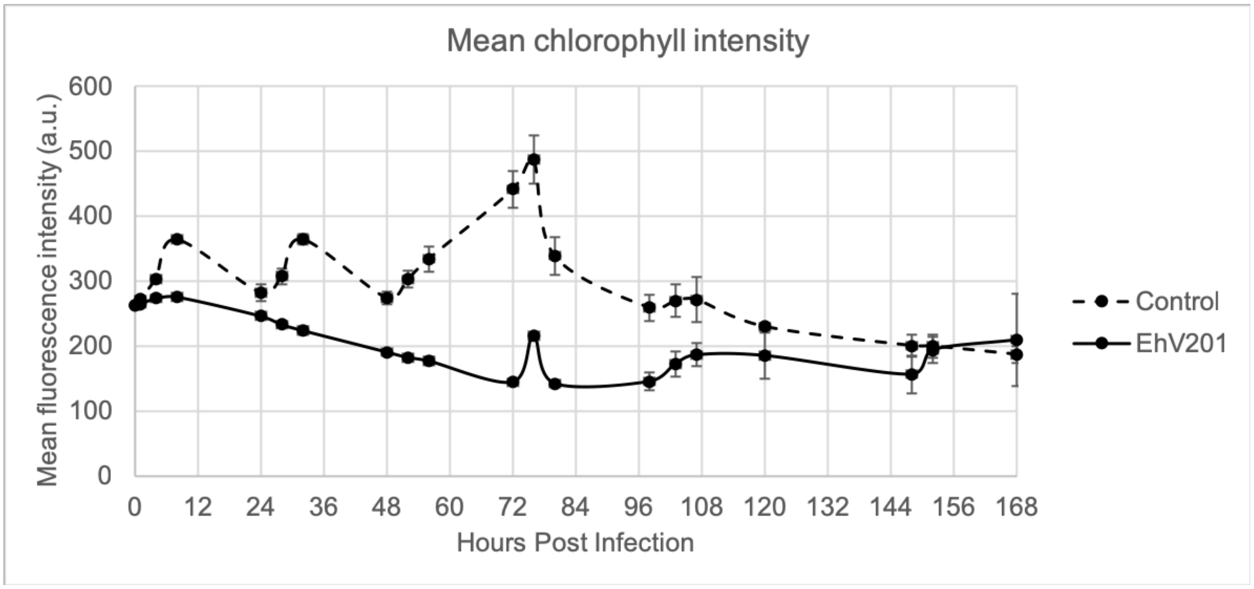


**Fig. S6. Chlorophyll mean intensity during infection.** Cell counts are based on chlorophyll fluorescence detected by flow cytometry. The cell gate selects cells that have high chlorophyll intensity (> $10^{\wedge 4}$ a.u). However, the mean chlorophyll intensity of that population can fluctuate as shown above between control (dashed line), and infected (full line) populations. Cells were infected at 9 a.m (0 hpi).

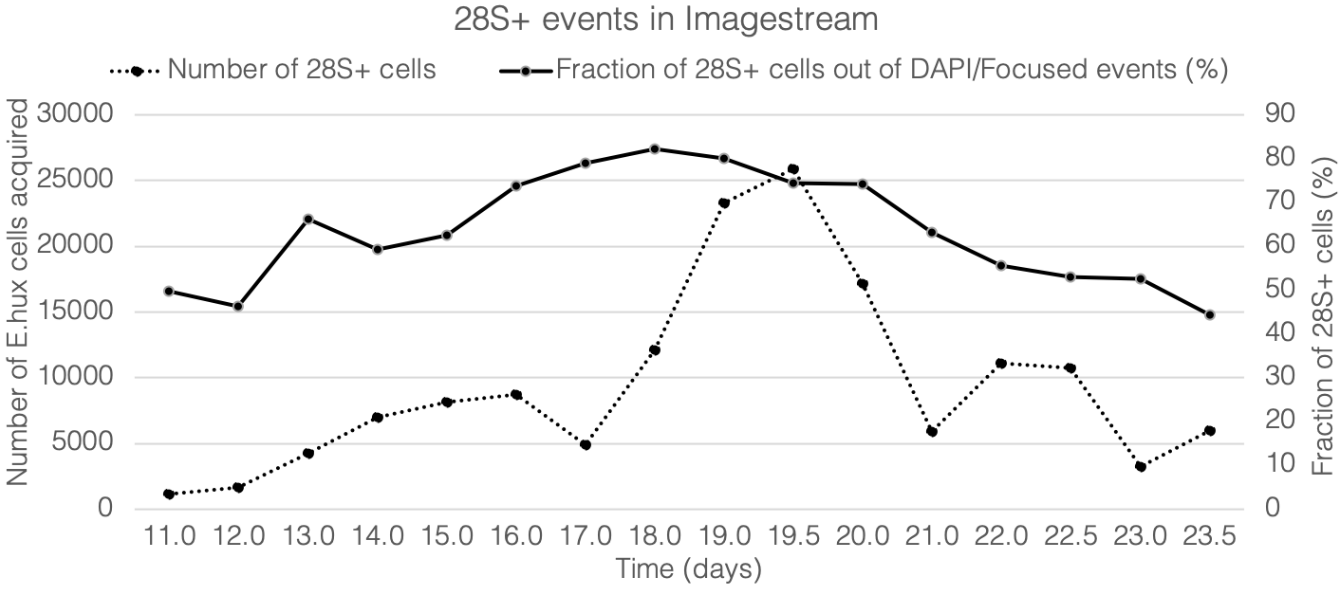


**Fig. S7. Adapting Virocell-FISH to specifically target *E. huxleyi* in environmental samples.** By designing a fluorescent probe that specifically targets the 28S ribosomal region of *E.huxleyi*, we were able to identify *E.huxleyi* cells in samples originating from a natural complex communities. Out of all DAPI+ events (cells with intensity value in the DAPI channel higher than $1.1*10^4$ a.u), the absolute number of *E.huxleyi* cells (left axis, dotted line) analyzed at each time point after day 16 was above 5000 cells, reaching 25000 single cells on day 19.5 (evening sample). The fraction of *E.huxleyi* cells (defined as 28S+ based on rRNA fluorescence, right axis, straight line) ranged between 40% at the beginning and end of the bloom to above 80% at the peak of the *E. huxleyi* bloom between days 17 and 19. DAPI+ events include all alga/grazers containing dsDNA that are below 40 micron in size.

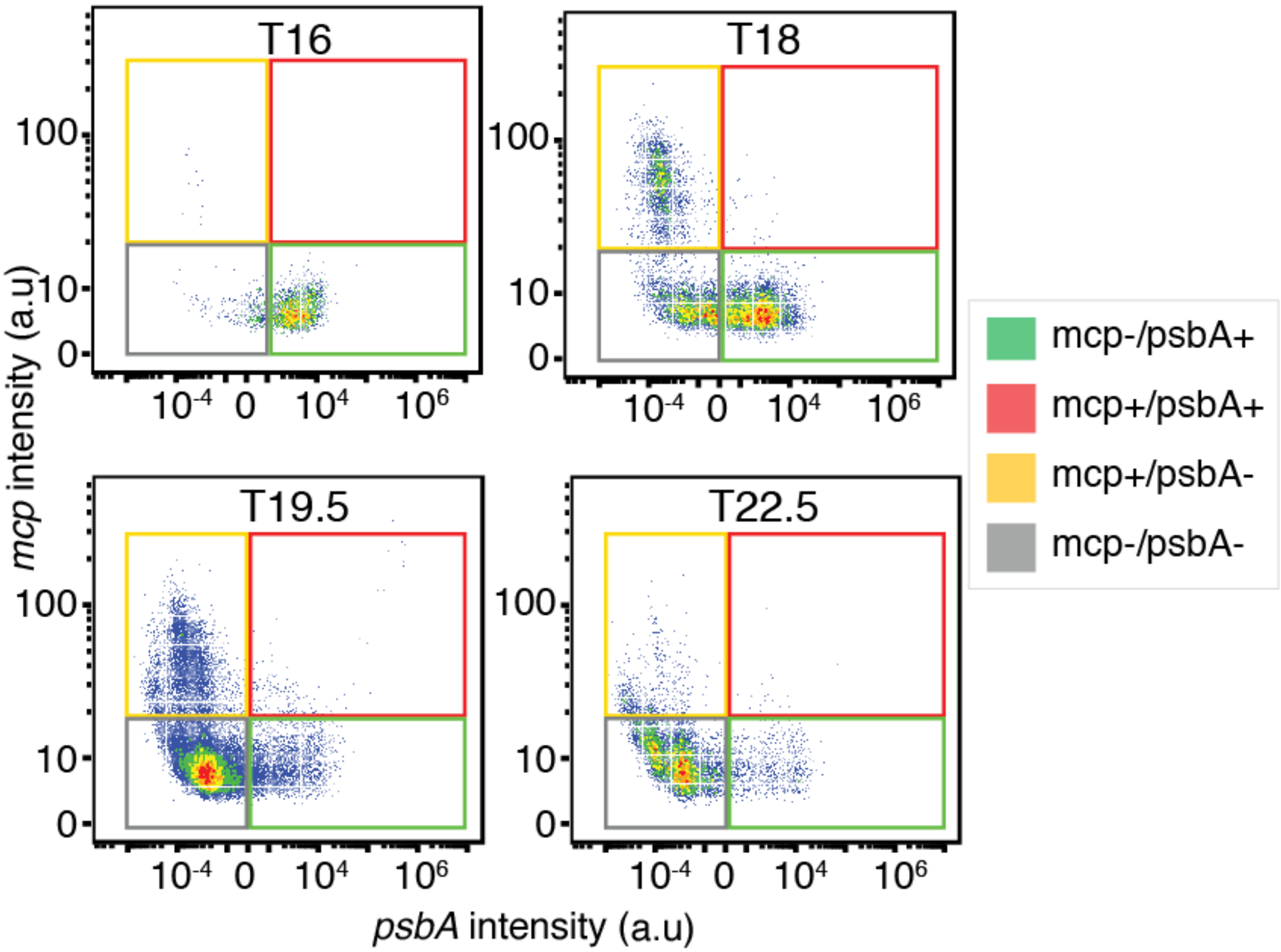


**Fig. S8. Defining *mcp*/*psbA* subpopulations in environmental samples.** To investigate *E. huxleyi* virocell heterogeneity during bloom succession, we plotted host and virus mRNA expression in parallel at different time points of days post infection (T16 to T22.5). The X axis represents the value of the probe intensity (in fluorescent arbitrary units) targeting the *psbA* host gene, and the Y axis represents the value of the probe intensity targeting the viral *mcp* gene. Using a threshold of 20 arbitrary units of max pixel in the *mcp* channel to define *mcp*+, cells and 0 arbitrary units of fluorescence intensity in the *psbA* channel to define *psbA*+ cells, we define four subpopulations as a combination of *mcp* and *psbA* signals.

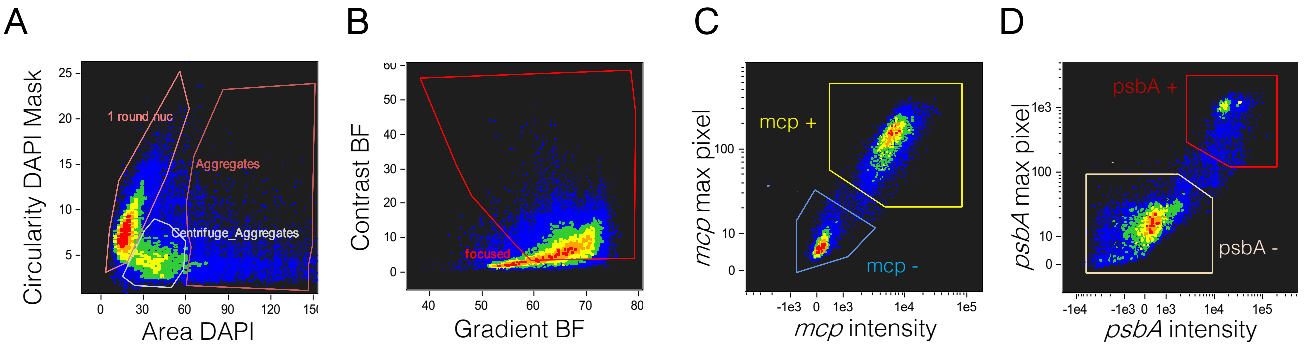


**Fig. S9. Pipeline of single cell identification in ISX.** (**A**) Identification of single cells based on the area of the DAPI in each event versus the circularity of the DAPI mask. (**B**) Identification of focused single cells based on the gradient in the brightfield versus the contrast. (**C**, **D**). Within the focused single cells, positively stained populations of *mcp* and *psbA,* respectively, can be defined by the intensity (sum of all pixel intensities in a single cell) versus the max pixel (value of the highest pixel) of each event.

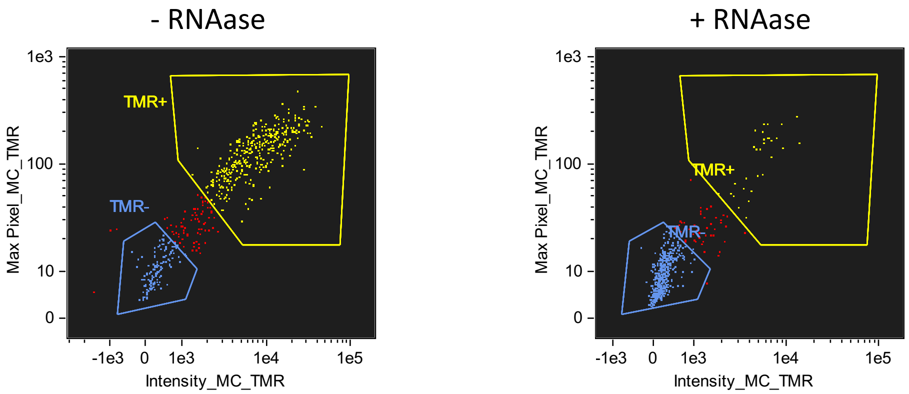


Intensity_MC_TMR, Max Pixel_MC_TMR

| Population | Count | %Gated |
|---|---|---|
| focused & DAPI+ & 1 nuc | 559 | 100 |
| TMR- & focused & DAPI+ & 1 nuc | 139 | 24.9 |
| TMR+ & focused & DAPI+ & 1 nuc | 344 | 61.5 |

Intensity_MC_TMR, Max Pixel_MC_TMR

| Population | Count | %Gated |
|---|---|---|
| focused & DAPI+ & 1 nuc | 594 | 100 |
| TMR- & focused & DAPI+ & 1 nuc | 517 | 87 |
| TMR+ & focused & DAPI+ & 1 nuc | 32 | 5.39 |

**Fig. S10. Comparison of the Virocell-FISH method with or without RNAase treatment.**

TMR is the abbreviation for "Tetramethylrhodamine" and is the fluorophore used to visualize the *mcp* transcripts. The non-treated sample shows 24.9% of *mcp*+ cells, whilst the RNAse treated sample showed less than 5% of *mcp*+ cells that could be noise considering the number of cells. This suggests that our Virocell-FISH probe catches mainly RNA and not DNA.

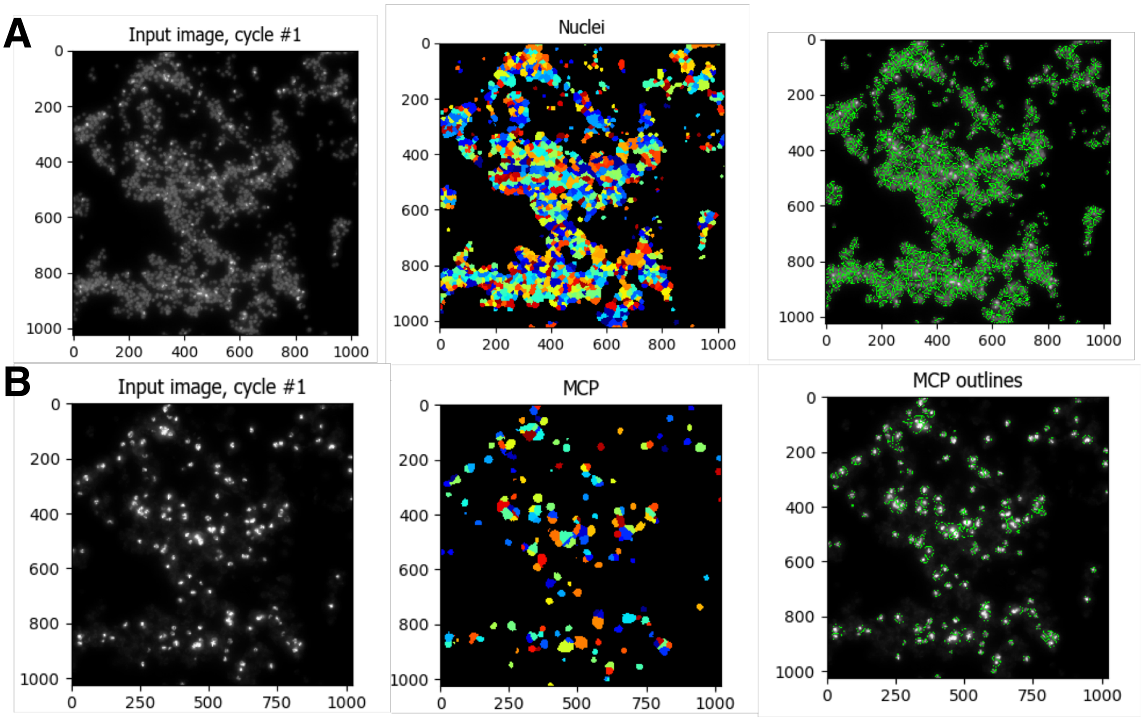


**Fig. S11. Quantifying infected cells in microscopy data using CellProfiler.** CellProfiler[3] was used to analyze microscopy data. In particular, the "PercentPositive" pipeline was applied. For each sample, several channels are collected including DAPI, TMR (*mcp* gene), Cy5 (*psbA* gene), and FITC (DioC6 membrane stain). (**A**) CellProfiler identifies cells based on the DAPI signal and draws cell contour. (**B**). Here, we show how the *mcp* images are then used to count how many cells are positive. A ratio of positive events is given as final output.

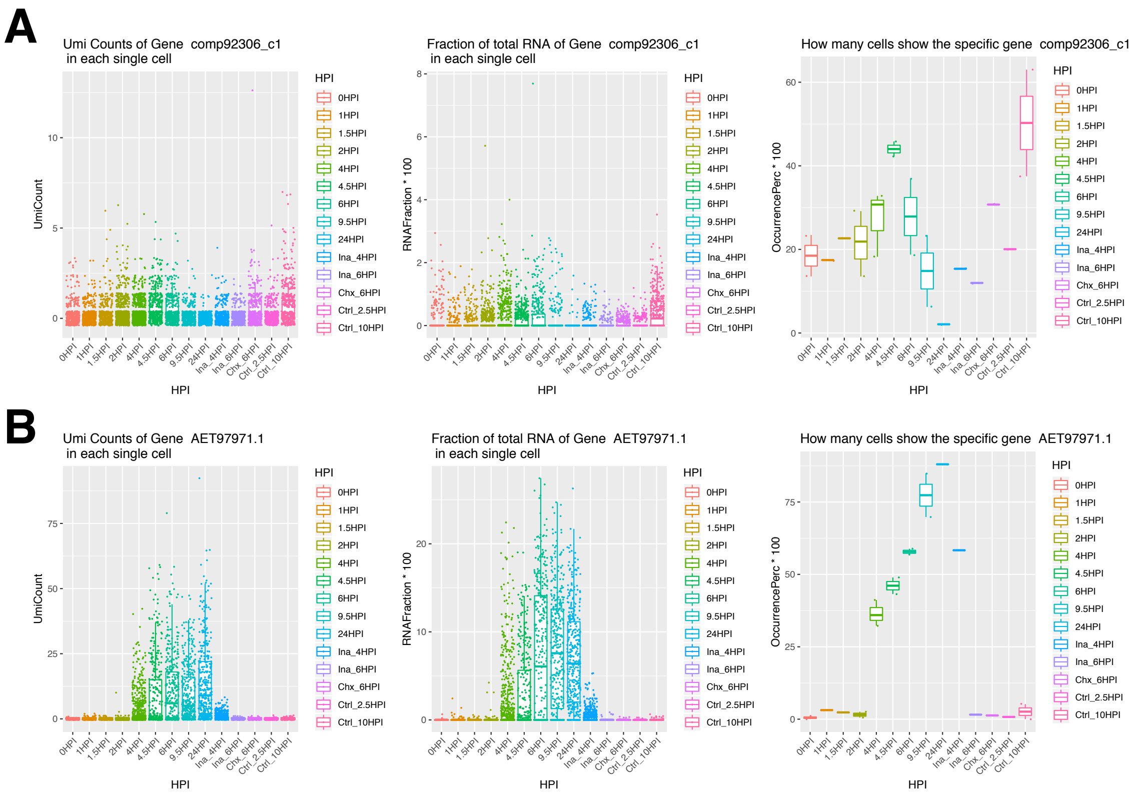


**Fig. S12. Gene expression of *psbA* and *mcp* in single cell RNA-seq sequencing data.** Expression of *psbA* and *mcp* in Single cell RNA-seq dataset during viral infection for a time course between 0-24 hpi for (**A**) *psbA* (**B**) *mcp*. This includes control, UV inactivated viruses, and cycloheximide experiments. From left to right: absolute unique molecular identifier (UMI) count per cell; ratio between gene UMI count and total UMI count in each cell; fraction of cells that show UMI for that gene. Data extracted from[4].

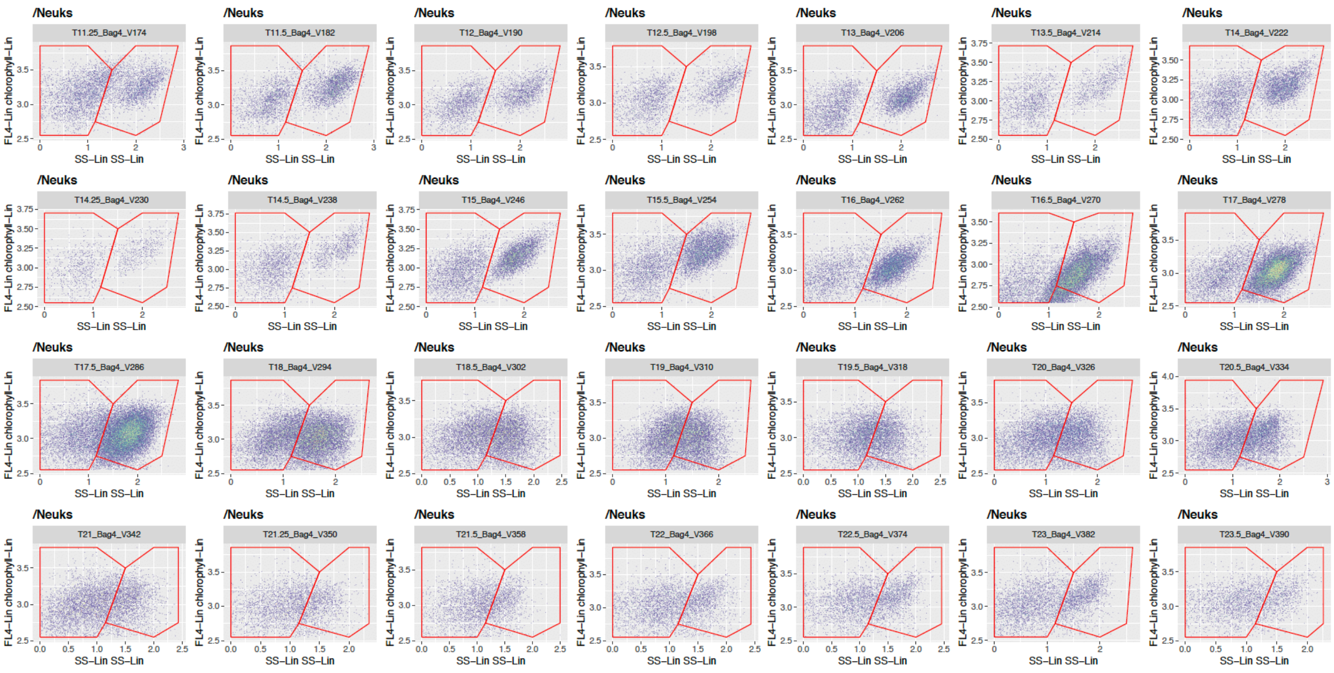


**Fig. S13. Flow cytometry gates used for *E. huxleyi* cells in the field.** Flow cytometry gates used for counting calcified and non-calcified nano-eukaryotes based on side scatter versus chlorophyll intensity shown here between day 11 and 23.5 which correspond to the bloom and demise phase. Absolute abundance of calcified *E.huxleyi* in cells/mL was obtained by normalizing to the sampled volume for each acquisition.

| | |
|---|---|
| MPN (infectious particles ml-1) | 7.6E+07 |
| E.huxleyi concentration (cells $ml^{-1}$) | 557000 |
| Volume of virus used (mL) | 38 |
| Exact MOI | 6.4 |

**Table S1: Calculation of virus:host ratio using the MPN method.** The concentration of infectious particles was calculated based on MPN method[6], performed on day 0 of the time course of infection, with the same cells and viral stock used for the experiment in **Fig. 1D**. The final virus:host ratio performed in the experiment was of 6.4
